# Axonal mitochondria across species adjust in diameter depending on thickness of surrounding myelin

**DOI:** 10.1101/850370

**Authors:** Benjamin V Ineichen, Keying Zhu, Karl E Carlström

## Abstract

In the central nervous system (CNS), axons and its surrounding myelin sheaths, generated by oligodendrocytes, greatly depend on each other, where oligodendrocytes provide axons with both trophic and metabolic support. Across spices, assessment of the axon-myelin ultrastructure is the key-approach to visualize de- and re-myelination of axons. However, this assessment omits to provide information on axonal homeostasis or how axon-myelin influence one another. Since mitochondria may adjust in size thus mirroring the intracellular physiological and metabolic status we applied this to myelinated axons in the CNS. We herein show that a large axonal mitochondria diameter correlates with thinner surrounding myelin sheaths across different CNS tracts and species, including human. We also show that the relation between axonal mitochondria diameter and surrounding myelin thickness is a valuable measurement to verify advanced remyelination in two commonly used experimental demyelinating models, namely the cuprizone and the lysolecithin (LPC) model. Lastly, we show that axonal mitochondria adjust in diameter in response to the thickness of the axonal surrounding myelin whereas the opposite adaption was absent. In summary, the link between axonal mitochondria diameter and surrounding myelin thickness provide insight on the axon-myelin relation both during homeostasis and pathological conditions. This link is also translational applicable and can thus contribute to a better understanding on how to study remyelination using experimental models.

## Introduction

The axon and the surrounding myelin sheath have a very close relationship, morphologically and functionally [11, 24]. This is e.g. illustrated by the association between axonal diameter and myelin thickness, that together influencing transduction in the central nervous system (CNS) [11, 13, 23]. During neuro-inflammatory and/or demyelinating insults, such as multiple sclerosis (MS), this relationship is disrupted. Although the coherence between the myelin and the axon can be re-established during subsequent remyelination, the regenerated myelin sheaths are thinner compared to pre-injury condition. Following insults to the myelin, adaptive mechanisms do occur in the axons as chronically demyelinated and remyelinated axons display higher mitochondria density compared to homeostatic conditions [1, 20, 29]. Similar, mitochondrial adaptation mechanisms have also been observed outside the CNS where mitochondria growth/enlargement is an accepted proxy for cellular adaption to increased energy demand and/or a detrimental response to cellular stress and Bax/Bcl [15, 21, 26]. Suggestively, the myelin sheath thickness and metabolic axonal features influence each other during myelin repair but also during homeostatic conditions [4, 5, 9]. However, there is an insufficient understanding as to which processes govern this interplay, more specific: which processes regulate the myelin sheath thickness and/or the axonal energy metabolism during homeostasis, but also during injury.

We serendipitously found in transmission electron microscopy (TEM) sections from naïve rat corpus callosum (CC), a strong correlation between myelin sheath thickness and axonal mitochondrial diameter. We thus further explored this association as a potential valuable measurement of the close interplay between myelin sheath thickness and axonal energy metabolism. This association was therefore further explored under physiological conditions and disrupted myelin homeostasis. We did this by re-analyzing both published and original TEM images from different species and CNS tracts.

## Materials and Methods

### TEM analysis

G-ratio and axonal mitochondria circumference (throughout referred to as axonal mitochondria diameter) from images unpublished or originally published in papers summarized in Table 1, were analyzed/re-analyzed using ImageJ with G-ratio plugin [12]. Axonal mitochondria display a limited variation in circumference, thus truncated axonal mitochondria caught in a TEM cross-section are a representative measurement for the whole organelle [19]. For all TEM images, all mitochondria-containing axons were analyzed, the mean mitochondria diameter were used as the representative value for every axon. To evaluate Plp.tg and Ax:Mfn2, additional images were obtained from the original authors. All statistical analyses were performed using GraphPad Prism Software. All correlation analyses were performed with Pearson’s *r* test. Multiple comparisons were throughout analyzed with one-way ANOVA with Bonferroni correction. *P*<0.05 was throughout considered as statistical significant.

## Results

### Myelin thickness correlates to axonal mitochondria diameter in rodents, macaque and man during homeostasis

Assessment of the axon-myelin ultrastructure using TEM and G-ratio is the golden standard for evaluation of homeostatic myelination and remyelination following injury [10]. G-ratio represents a relative measure for myelin thickness where a G-ratio of 1 represents a completely unmyelinated/naked axon and G-ratio <1 represent increasing myelin sheath thickness (Fig.1A). TEM also offers the possibility to assess additional ultra-structures, including axonal mitochondria, which are readily identified (Fig.1B).

**Fig.1.**
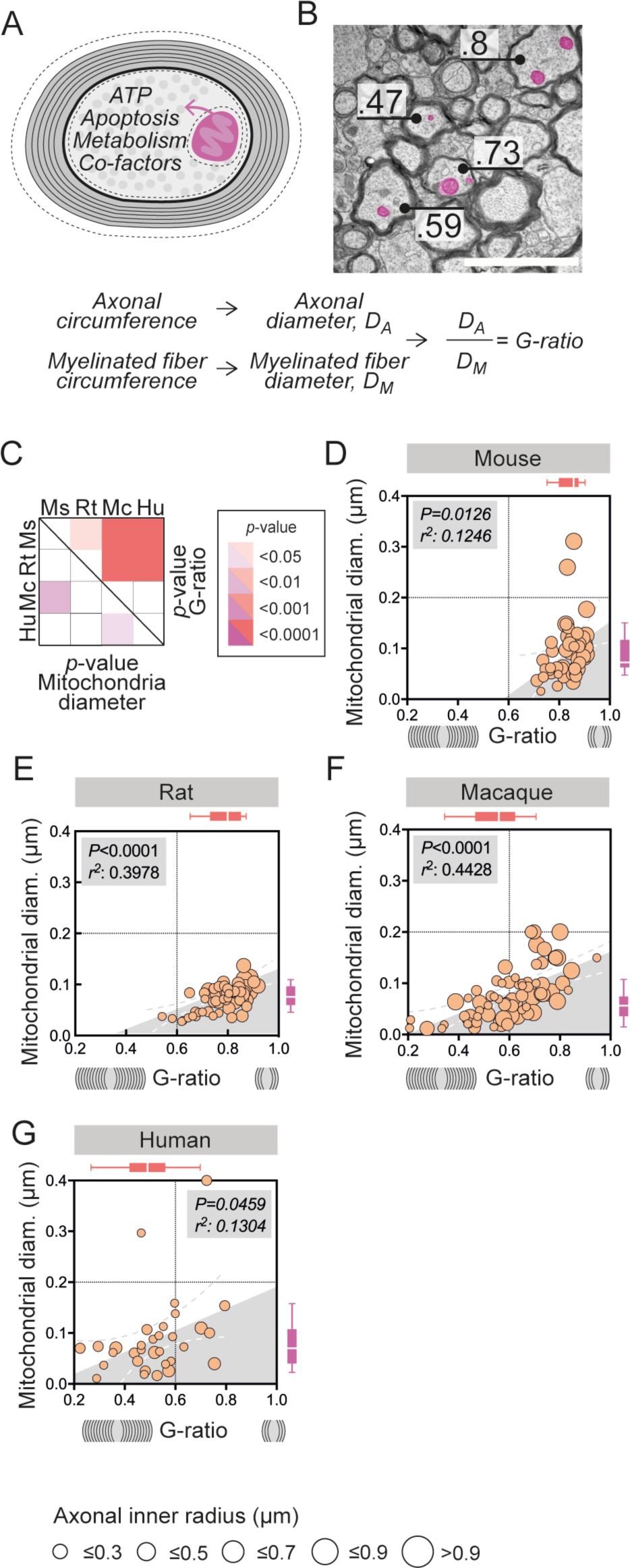
Myelin thickness correlates to axonal mitochondria diameter in rodents and macaque during homeostasis. (a) Illustration of a cross-sectional myelinated axon with the G-ratio representing the axonal diameter in relation to the diameter of both the axon and the surrounding myelin. (b) Representative TEM image of the rat corpus callosum (CC) with annotated G-ratio and pseudo-colored axonal mitochondria in purple (Scale-bar 1μm). (c) Summary of p-values comparing homeostatic G-ratio (red gradient) and mitochondria diameter (purple gradient) between all species (mouse (Ms), rat (Rt), macaque (Mc) and human (Hu). (d) Homeostatic mitochondria diameter (μm) and G-ratio in mouse CC (n_animals_=1 n_total axons_ =49). (e) Homeostatic mitochondria diameter (μm) and G-ratio in rat CC (n_animals_=2, n_total axons_ =61) (f) Homeostatic mitochondria diameter (μm) and G-ratio in macaque spinal cord (SC) (n_animals_=3, n_total axons_=72). (g) Homeostatic mitochondria diameter (μm) and G-ratio in man prefrontal cortex (n_subjects_=3, n_total axons_=32). The diameter of each point of measure indicates the axonal inner radius in μm. All correlation analyses were performed with Pearson’s r test. Multiple comparisons in (c) were performed using one-way ANOVA.

During homeostatic conditions, G-ratio and mitochondria diameter varied among the assessed species (Fig.1C). However, the G-ratio and mitochondria diameter correlated in rodents, where CC and spinal cord (SC) axons with thinner myelin sheaths had larger axonal mitochondria (Fig.1D-E, Table I). The correlation between myelin sheaths and axonal mitochondria diameter was further conserved in higher species, demonstrated by a similar correlation in macaque SC (Fig.1F, Table I) [25]. Finally, this was also verified in human post-mortem tissue (Fig.1G, Table I) [18, 27]. No differences were found in numbers of mitochondria in any of the assessed species or CNS regions (data not shown).

### Association between myelin thickness and axonal mitochondria diameter during de- and remyelination

Demyelination and sequent remyelination are major and metabolic stressful events for the axon. We thus assessed whether myelin-axons were able to maintain the positive correlation between axonal mitochondria diameter and G-ratio during de- and remyelination. The two commonly used experimental models to study demyelination and subsequent remyelination uses; cuprizone, a dietary copper-chelating agent, and lysolecithin (LPC), an intraparenchymally injected detergent [3, 14, 30].

In LPC-lesioned mice, the G-ratio was increased during early remyelination (d10 post injection) compared to advanced remyelination (d24 post injection). A similar decrease in G-ratio was observed during demyelination (5weeks) compared to early remyelination (6weeks) in cuprizone-fed mice (Fig.2A). Concomitantly, the axonal mitochondria diameter followed the same decrease when comparing the early and later time points (Fig.2B).

The correlation observed during homeostatic conditions was lost during demyelination in both models (Fig.2C left, D left). However, the correlation was re-established at later time-points, i.e. advanced remyelination in LPC and early remyelination in cuprizone (Fig. 2C right, D right).

**Fig.2.**
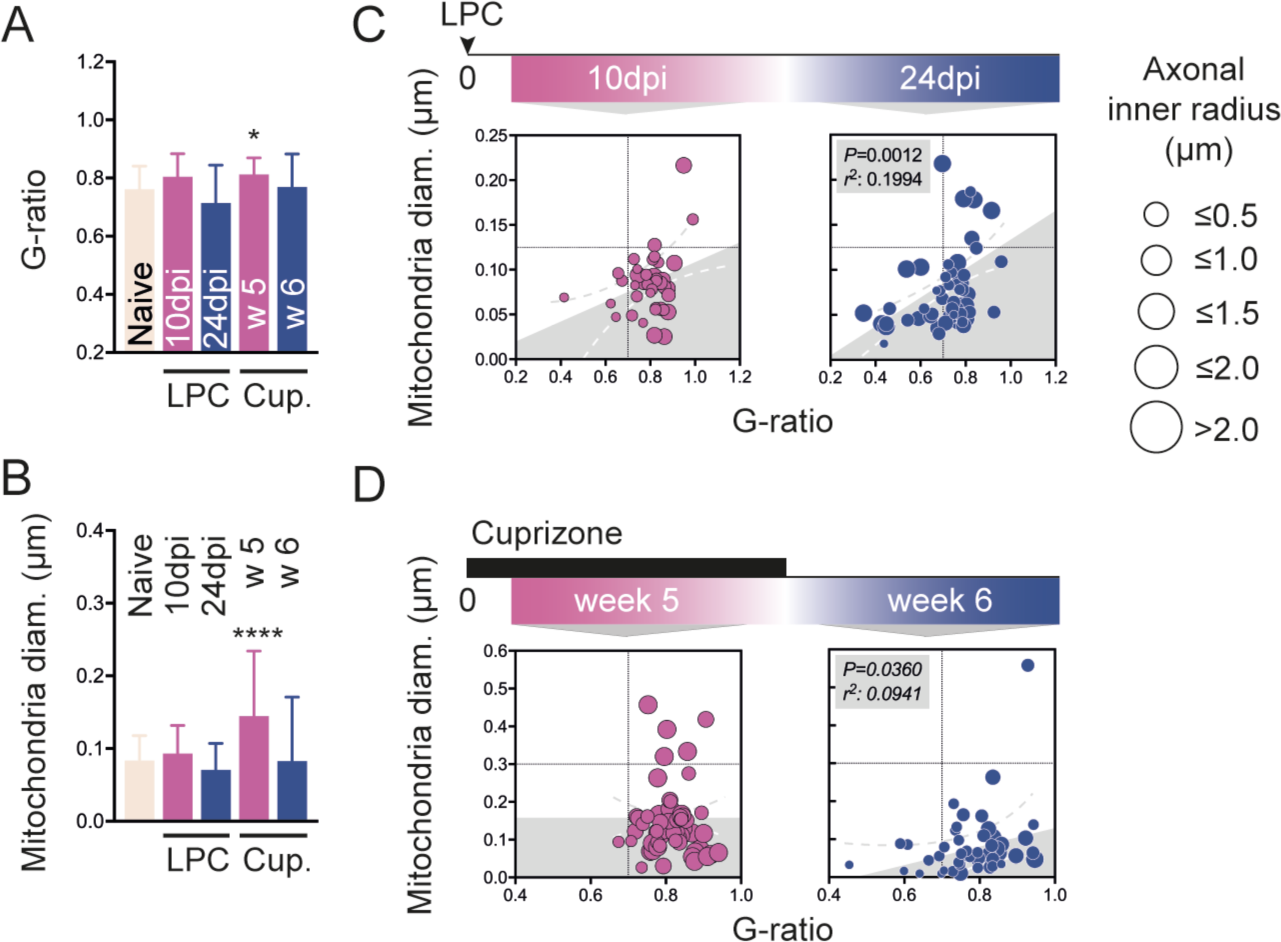
Axonal mitochondria diameter correlates with myelin sheath thickness during advanced but not early de- or remyelination. (a) G-ratio and (b) axonal mitochondria diameter in LPC and cuprizone model of toxic demyelination. Statistical comparison to naïve condition. (c) G-ratio plotted against axonal mitochondria diameter (μm) in rat CC (c left) 10 days following LPC injection (early remyelination, n_animals_=2, n_total axons_=39, p=0.0565) and (c right) 24 days following LPC injection (advanced remyelination, n_animals_=3, n_total axons_ =55). (d) G-ratio plotted against axonal mitochondria diameter (μm) in mice CC (d left) during cuprizone feeding (n_animals_=1, n_total axons_ =60) and a (d right) 1 week after cuprizone withdrawal (n_animals_=2, n_total axons_ =48). The diameter of each point of measure indicates the axonal inner radius in μm. Multiple comparisons to naïve condition in (a, b) were performed using one-way ANOVA. All correlation analyses were performed with Pearson’s r test. *p<0.05, ****p<0.0001

### Axonal mitochondria adapt the diameter to dysregulated myelination but not vice versa

Our data indicates a close correlation between G-ratio and axonal mitochondrial diameter during homeostasis and during advanced remyelination. But the direction of causality remains to be identified. Thus, to address whether increased axonal mitochondrial diameter is a consequence of thinner surrounding myelin or if axonal mitochondrial diameter affects the G-ratio, TEM images from strains with either dysfunctional myelin sheath composition or dysfunctional mitochondria function were assessed (Table I).

Analysis of axonal mitochondria and G-ratio of wild-type/heterogeneous mice of either genotype with normal myelin ultrastructure confirmed previously observed pattern where larger axonal mitochondria diameter correlated with larger G-ratio (Fig.3A, E). The correlation was significant for every individual strain as well as when pooled together (Fig.3A, E).

**Fig.3.**
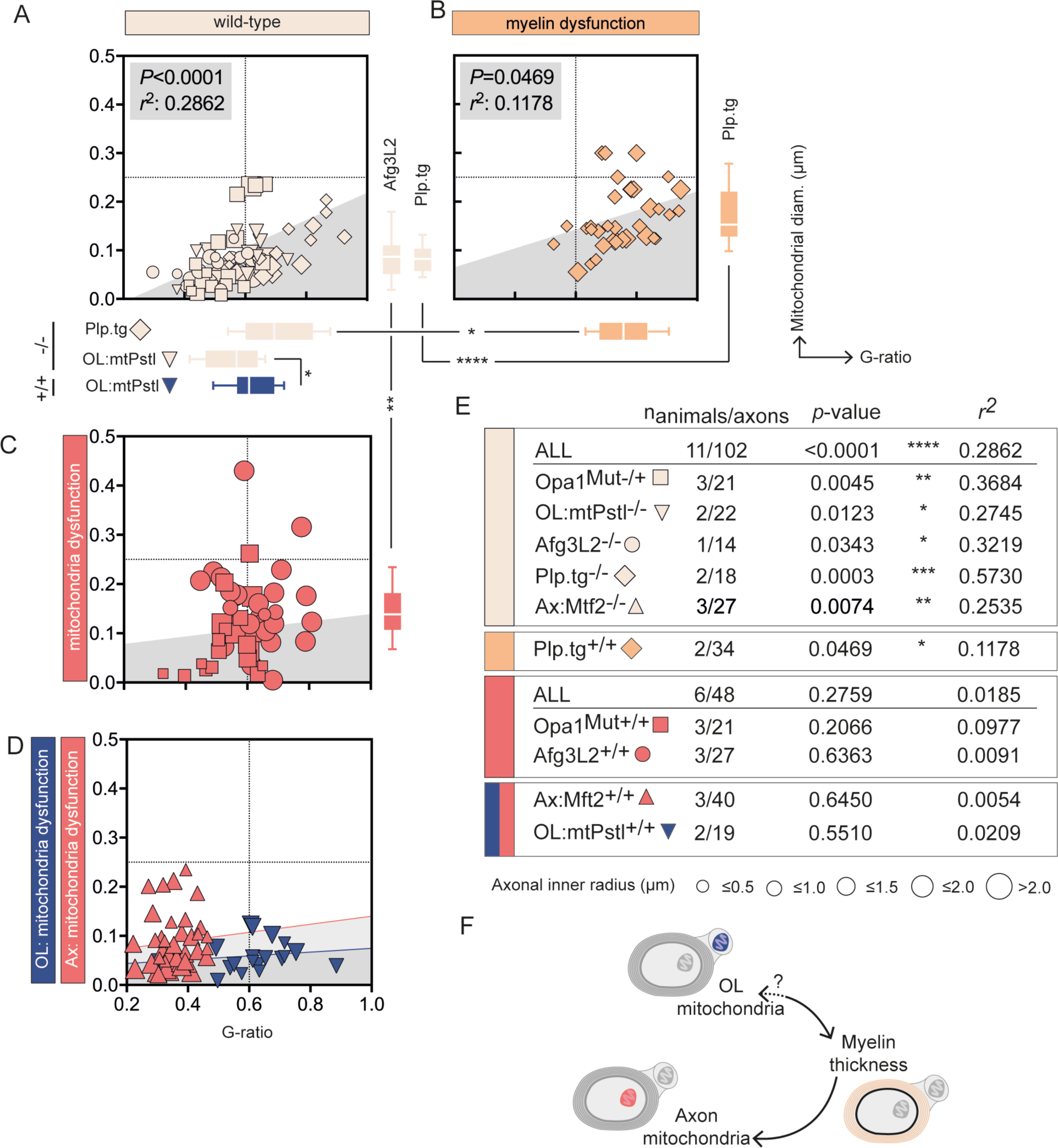
Hypo-myelination affects axonal mitochondria diameter but not vice versa. (a) Homeostatic mitochondria diameter and G-ratio in wild-type mice. (b) Homeostatic mitochondria diameter (μm) and G-ratio in Plp.tg^+/+^ mice (n_animals_=2, n_axons_=34). (c) Homeostatic mitochondria diameter (μm) and G-ratio in Opa1^Mut +/+^ (n_animals_=3, n_axons_=21) and Opa1^Mut +/+^ (n_animals_=3, n_axons_=27) mice. (d) Homeostatic mitochondria diameter (μm) and G-ratio in Ax:Mfn2^+/+^ (n_animals_=3, n_axons_=40) and OL:mtPstI^+/+^ (n_animals_=2, n_axons_=19) mice. and Opa1^Mut +/+^. (e) Statistical summary of correlation between G-ratio and axonal mitochondria diameter in all strains. (f) Schematic illustration of the adaption hierarchy where OL mitochondria dysfunction influence G-ratio (i.e. myelin thickness) which in turn influence axonal mitochondria diameter. All correlation analyses were performed with Pearson’s r test, multiple comparisons of mitochondria diameter and G-ratio in (a-c) were performed using one-way ANOVA. *p<0.05, **p<0.01, ****p<0.0001

The Plp.tg^+/+^ mice display dysfunctional myelin composition and hypo-myelination due to knock-in of myelin proteolipid protein (Plp) [16]. Indeed, this was confirmed by an increased G-ratio in Plp.tg^+/+^ compared to wild-type (Plp.tg^-/-^) (Fig.3A, B, Table II). Further, in response to the hypo-myelinated axons, axonal mitochondria increased in diameter (Fig.3A, B). Due to this mitochondrial adaption, Plp.tg^+/+^ mice maintain the correlation observed in wild-type conditions (Fig.1, Fig3A, B, E).

To address the opposite, i.e. whether mitochondria dysfunction would influence myelin thickness, we analyzed axonal mitochondria and G-ratios in mice with dysfunctional mitochondria. Mice with Opa^Mut+/+^ or Afg3L2^+/+^ mutations generates dysfunctional mitochondria as described previously [6, 22, 28]. Analysis of TEM images from Opa^Mut+/+^ or Afg3L2^+/+^ mice displayed increased mitochondria diameter compared to corresponding wild-types in both strains (Opa^Mut-/+^ or Afg3L2^-/-^); however, the difference was only statistically significant in Afg3L2^+/+^ mice compared to their wild-type, but not in Opa^Mut^ (*p*=0.1658) (Fig.3A, C). Intriguingly, neither the Opa^Mut+/+^ or Afg3L2^+/+^ strain showed lowered G-ratios (Fig.3A, C) and no correlation between G-ratio and axonal mitochondria diameter was seen in these strains with mitochondrial dysfunction (Fig.3C, E).

Opa^Mut+/+^ and Afg3L2^+/+^ are full-body mutants, thus mitochondria function is presumably altered in all cells including myelin-generating oligodendrocytes. Thus, mitochondrial dysfunction in oligodendrocytes could be responsible for the observed lack of G-ratio adaption to axonal mitochondria dysfunction. To address this, mice with axonal-specific mutations in the mitochondria protein Mtf2 (here referred to as Ax:Mfn2) were assessed [2]. However, similar to the full-body mutated strains (Opa^Mut+/+^ and Afg3L2^+/+^) the axon-surrounding myelin failed to adapt in the Ax:Mfn2^+/+^ in response to dysfunctional axonal mitochondria, thus leaving the G-ratio unchanged compared to wild-type (Ax:Mfn2^-/-^) (Fig.3A, D, E).

To verify whether also oligodendrocyte-specific loss of mitochondria function could interfere with G-ratio, mice expressing mtPstI under Plp promoter (here referred to as OL:mtPstI) were analyzed. OL:mtPstI, causing mitochondria DNA-breakage, rendered a higher G-ratio (i.e a thinner myelin sheath) (Fig.3A, D). However, OL:mtPstI did not indicate any implication on axonal mitochondria diameter from dysfunctional oligodendrocyte mitochondria (*p*=0.1430) [28]. Also, the correlation between mitochondria diameter and G-ratio were lost (Fig.3E). Taken together, our data support the hypothesis that axonal mitochondria can adapt to loss of myelin integrity. However, oligodendrocytes seem not to adjust G-ratio in a state of dysfunctional axonal mitochondria.

## Discussion

The axon and its surrounding myelin have a close relationship, yet fundamental understanding of their mutual regulation processes is lacking. We herein explored the interplay between G-ratio (i.e. myelin sheath thickness) and axonal mitochondrial diameter, i.e. the thinner the myelin sheath surrounding an axon, the larger the corresponding axonal diameter.

Interestingly, we could confirm a strong correlation in different spinal cord and brain tracts as well as across spices, but the correlation was lost during de- and/or early remyelination. Finally, data from transgenic mice with either dysfunctional myelin or mitochondria suggest that mitochondria adjust to the G-ratio but not *vice versa*. The main restriction of our study is the potential selection bias caused by partial analysis of printed publication figures. However, main findings were confirmed in additional original material. Moreover, the findings were robust when investigating different species, mutant strains and CNS regions.

There is a strong correlation between G-ratio and axonal mitochondria diameter in all assessed species; rodents, non-human primates and humans. Interestingly, current literature describes discrepancies in the contribution from non-dividing oligodendrocytes during remyelination and axonal protection depending on the assessed specie [7, 8]. In that perspective, we herein identify a conserved feature, which can further help us understand differences and similarities in axon-myelin status between conditions and species.

Further, we herein show that the correlation between thinner myelin sheets and larger axonal mitochondria is re-established during later stages of remyelination in two commonly used models of toxic demyelination. In the early remyelination phase (ten days post LPC injection) or during peak demyelination phase (after 5 weeks cuprizone administration) mitochondrial diameter and G-ratio were not correlated. The correlation was, however, re-established upon remyelination (24 days post LPC injection or 1 week after withdrawal of cuprizone administration). The lack of this correlation during demyelination or early remyelination may indicate ongoing axonal adaptation processes. Assessment of G-ratio in correlation to mitochondria diameter thus indicates a valuable read-out to identify advanced remyelination and re-establishment of homeostatic relation between axon and surrounding myelin. Compared to current measurement, this better reflect axonal energy balance and could thus be used in addition to traditional read-outs when evaluating remyelination.

The lack of mitochondrial adaption immediately following LPC or cuprizone but observed upon genetic disruption of myelin in the Plp.tg^+/+^ indicates that the mitochondrial adaption is not instant upon loss of myelin. This further underlines the usability of the mitochondria-myelin correlation to better determine fulfilled remyelination with established metabolic function. To address direction of causality between G-ratio and axonal mitochondria diameter, different transgenic strains with either dysfunctional myelin or mitochondria were re-analyzed. Interestingly, axonal mitochondria in the hypo-myelinated Plp.tg^+/+^ adapted to the thin myelin and increased in diameter, thus Plp.tg^+/+^ maintained the correlation but with larger G-ratio and mitochondria diameter as observed in Plp.tg^-/-^. The increase in mitochondrial diameter might be an axonal adjustment in order to compensate for insufficient axonal energy supply from the myelin sheath and/or compensation for the lack of sufficient myelination to maintain function [11]. In addition, our findings underline the role of oligodendrocytes as providers of metabolic support to insulated axons [11, 17].

Our data indicate further that the G-ratio seems not to adjust to dysfunctional mitochondria. Independent of the nature of mitochondria dysfunction, the surrounding myelin sheets were incapable to adjust its thickness. Together, this indicates that thinner myelin sheaths induce adjustment in axonal mitochondria diameter. But the axon-surrounding myelin is not able to adapt to loss of mitochondria function potentially causing axonal energy depletion and loss of axonal velocity.

## Conclusions

Our data from different species and CNS regions indicate that axonal mitochondria seem to adjust their diameter depending on the thickness of the axon-surrounding myelin sheaths. This further underlines a close and dynamic interplay between axon and myelin. This improved understanding can contribute to better understanding on how to study remyelination in experimental models. Also, to pave the way for the development of remyelinating and/or neuroprotective therapies in demyelination disorders such as multiple sclerosis.

## List of abbreviations

SC: spinal cord
CC: corpus callosum
LPC: lysophosphatidylcholine
CNS: central nervous system
TEM: transmission electron microscopy
MBP: myelin basic protein

## Author contribution

KC, BI and KZ conceived the study and wrote the manuscript.

## Acknowledgement

We acknowledge Dr. Mark McLaughlin; University of Glasgow, Dr. Nathalie Bernard-Marissal; École Polytechnique Fédérale de Lausanne, Dr, Roman Chrast; Karolinska Institute and Dr. Jun Wang, Fudan University for kindly providing TEM images.

