## Supplementary material for "Axonal mitochondria across species adjust in diameter depending on thickness of surrounding myelin": Table I

### Table I: Summary strains

| Species | Strain | Cell-specificity | Phenotype | CNS region | Condition | Figure | Ref |
|---|---|---|---|---|---|---|---|
| Mouse | C57BL/6 | N/A | N/A | CC | Naive | 1D | |
| Rat | DA | N/A | N/A | CC | Naive | 1E | Carlstrom et al 2019 (submitted manuscript) |
| Macaque | N/A | N/A | N/A | SC | Naive | 1F | Stikov et al., 2015 |
| Human | N/A | N/A | N/A | Prefrontal cortex | Bipolar disorder | 1G | Uranov et al., 2001, Lewis et al., 2019 |
| Rat | DA | N/A | N/A | CC | 10d post LPC | 2 | Carlstrom et al 2019 (submitted manuscript) |
| Rat | Lewis | N/A | N/A | SC (dorsal funiculus) | 24d post LPC | 2 | Ineichen et al., 2017 |
| Mouse | C57BL/6 | N/A | N/A | CC | 5w of cuprizone | 2 | |
| Mouse | C57BL/6 | N/A | N/A | CC | 1w post cuprizone withdrawl | 2 | |
| Mouse | Plp.tg (C57BL/6J) | No | Hypo-myelinated axons | SC (cervical) | Naïve: Plp.tg-/-, Plp.tg+/+ | 3 | Karim et al. 2007 |
| Mouse | Opa1Mut (C57BL/6J) | No | Dysfunctional mitochondria | Optic nerve | Naïve Opa1Mut-/+, Opa1Mut+/+ | 3 | Chao de la Barca et al., 2017, Davies et al., 2007 |
| Mouse | OL:mtPstI (C57BL/6) | Oligodendrocyte (Plp-Cre) | Mitochondrial DNA break in OL | SC (thoracic) | Naïve: OL:mtPstI-/-, OL:mtPstI+/+ | 3 | Madsen et al., 2012 |
| Mouse | Afg3L2 (C57BL/6N) | No | Dysfunctional mitochondria | SC (lumbar) | Naïve: Afg3L2-/-, Afg3L2+/+ | 3 | Wang et al., 2016 |
| Mouse | Ax:Mfn2 (C57BL/6) | Neurons (Eno2) | Dysfunctional mitochondria in neurons | SC (lumbar) | Naïve: Ax:MFNR94Q, Ax:MFNWT | 3 | Bernard-Marissal et al., 2019, Cartoni et al., 2010 |

*Abbreviations: CC, corpus callosum; DA, Dark agouti; SC, spinal cord; OL, oligodendrocyte*
