## Supplementary material for "Axonal mitochondria across species adjust in diameter depending on thickness of surrounding myelin": Table II

### Table II: Summary statistics

| | Mitochondria diam. | *p* -value' | G-ratio | *p* -value' |
|---|---|---|---|---|
| Opa1Mut-/+ | 0.059±0.036 | | 0.536±0.101 | |
| Opa1Mut+/+ | 0.085±0.069 | | 0.535±0.085 | |
| OL:mtPstI-/- | 0.057±0.025 | | 0.549±0.096 | * |
| OL:mtPstI+/+ | 0.048±0.019 | | 0.622±0.076 | |
| Afg3L2-/- | 0.086±0.054 | ** | 0.590±0.084 | |
| Afg3L2+/+ | 0.155±0.083 | | 0.634±0.093 | |
| Plp.tg-/- | 0.081±0.030 | **** | 0.700±0.126 | * |
| Plp.tg+/+ | 0.168±0.064 | | 0.751±0.105 | |
| Ax:Mtf2-/- | 0.069±0.027 | | 0.364±0.074 | |
| Ax:Mtf2+/+ | 0.085±0.056 | | 0.348±0.072 | |

'means compared with one-way ANOVA
